# Removal of the C6 vaccinia virus interferon-β inhibitor in the hepatitis C vaccine candidate MVA-HCV elicited in mice high immunogenicity in spite of reduced host gene expression

**DOI:** 10.1101/330902

**Authors:** María Q. Marín, Patricia Pérez, Carmen E. Gómez, Carlos Óscar S. Sorzano, Mariano Esteban, Juan García-Arriaza

**Affiliations:** Department of Molecular and Cellular Biology, Centro Nacional de Biotecnología (CNB), Consejo Superior de Investigaciones Científicas (CSIC), Madrid, Spain.; Biocomputing unit, Centro Nacional de Biotecnología (CNB), Consejo Superior de Investigaciones Científicas (CSIC), Madrid, Spain.

**Keywords:** HCV, poxvirus, MVA, vaccine, C6L, interferon, host gene expression, mice, cellular responses, humoral responses

## Abstract

Hepatitis C virus (HCV) represents a major global health problem for which a vaccine is not available. MVA-HCV is a unique HCV vaccine candidate based in the modified vaccinia virus Ankara (MVA) vector expressing the nearly full-length genome of HCV genotype 1a that elicits broad and polyfunctional CD8^+^ T-cell responses in mice. With the aim to improve the immune response of MVA-HCV and due to the importance of interferon (IFN) in HCV infection, we deleted in MVA-HCV the vaccinia virus (VACV) *C6L* gene, encoding an inhibitor of IFN-β that prevents activation of the transcription factors IRF3 and IRF7. The resulting vaccine candidate (MVA-HCV ΔC6L) expresses all HCV antigens and deletion of *C6L* had no effect on viral growth in permissive chicken cells. In human monocyte-derived dendritic cells, infection with MVA-HCV ΔC6L triggered severe down-regulation of IFN-β, IFN-β-induced genes and cytokines similarly to MVA-HCV, as defined by real-time PCR and microarray analysis. In infected mice both vectors had a similar profile of recruited immune cells and induced comparable levels of adaptive and memory HCV-specific CD8^+^ T-cells, mainly against p7+NS2 and NS3 HCV proteins, with a T cell effector memory (TEM) phenotype. Furthermore, antibodies against E2 were also induced. Overall, our findings showed that while these vectors had a profound inhibitory effect on gene expression of the host, they strongly elicited CD8^+^ T cell and humoral responses against HCV antigens. These observations add support to the consideration of these vectors as potential vaccine candidates against HCV.

**IMPORTANCE:** Hepatitis C virus represents a global health problem with 71 million of people infected worldwide. While direct-acting antivirals agents can cure hepatitis C virus infection in most of patients, their high cost and the emergence of drug resistant variants make them not a feasible and affordable strategy to eradicate the virus. Therefore, a vaccine is an urgent goal that requires efforts in understanding the correlates of protection for hepatitis C virus clearance. Poxvirus vectors, in particular the attenuated modified vaccinia virus Ankara, are ideal as vaccine candidates due to their ability to induce both T and B cell immune responses against heterologous antigens and protection against a wide spectrum of pathogens. Here we describe the generation, genetics and immunogenicity elicited by MVA-HCV ΔC6L, a novel vaccine candidate for hepatitis C virus that expresses nearly all of hepatitis C proteins but lacks an IFN-β inhibitor, the C6 vaccinia virus protein.

## INTRODUCTION

Hepatitis C virus (HCV), a member of the genus *Hepacivirus*, family *Flaviviridae*, is an enveloped icosahedral virus, with positive-sensed and single-stranded RNA genome of 9,600 nucleotide bases long (1). HCV infection is a global health problem for which no vaccine is available. Upon HCV infection, 15-30% of people are able to clear the virus while the remaining 70-85% will develop a chronic infection that can end with liver cirrhosis and hepatocellular carcinoma. It is estimated that 71 million people suffer from chronic hepatitis C and approximately 400,000 die every year due to HCV-related diseases, as reported by the World Health Organization (http://www.who.int/mediacentre/factsheets/fs164/en).

Recently approved direct-acting antiviral agents (DAAs) are able to cure HCV at high rates, but they still have important limitations such as the emergence of drug resistance variants, their high cost, safety issues especially in advanced chronic disease, viral genotype dependency, and the fact that most individuals who are chronically infected are unaware of their infection and therefore unlikely to seek treatment (2, 3). Moreover, DAAs do not prevent future infections (4), hence the development of a prophylactic or therapeutic vaccine is a public health priority. Viral genome variability and immune evasion, lack of a suitable animal model and unclear correlates of protection during HCV infection represent the main hurdles for the development of a successful vaccine (5). However, spontaneous clearance of HCV infection and the ability to resolve subsequent infections in humans (6, 7) and chimpanzees (8) yield important evidence of a protective immune response with a memory component, making a vaccine a feasible goal.

Current efforts in HCV vaccine development are aimed to induce strong T cell and humoral immune responses that are able to target the most conserved viral antigens without producing liver immunopathology. It has been reported the importance of HCV-specific CD4^+^ and CD8^+^ T cell responses in clearance of primary infection and reinfection (9–11). Furthermore, recent studies have also proved the key role of antibody response to E1 and E2 HCV glycoproteins in the resolution of HCV infection (reviews in (11, 12). To date, several vaccine candidates against HCV have been tested with different rate of success (13). Among them, poxvirus vectors and in particular the modified vaccinia virus Ankara (MVA), possess ideal properties as vaccines due to their ability to induce strong T and B immune responses, their safety profile, low cost, ease of manufacture and administration and the low prevalence of anti-vector immunity in the global population (14–17).

However, more efficient and optimized vaccine candidates able to enhance both cellular and humoral immune responses against HCV antigens are desirable. One of the strategies is to enhance the induction of interferon (IFN), essential to trigger a potent defense against intracellular pathogens, such as HCV, and create an antiviral state in surrounding cells that will help to avoid chronic phase of infection (18, 19). Additionally, one of the main approaches that has been used to improve the immunogenicity of MVA recombinants is the deletion of immunosuppressive genes that are still present in the MVA genome (20). One of these vaccinia virus (VACV) immunomodulatory genes is *C6L*, encoding the C6 protein, a non-essential virulent factor expressed early in infection (21, 22). C6 interacts with the scaffold proteins TANK, NAP1 and SINTBAD and inhibits the kinases TBK1 and IKKε, which are the activators of the transcription factors IRF3 and IRF7, resulting in a blockade of IFN-β expression (21). Moreover, C6 inhibits type I IFN signaling in the nucleus and binds to the transactivation domain of STAT2 (23). Deletion of VACV *C6L* gene in the HIV/AIDS vaccine candidate MVA-B enhanced HIV-1-specific cellular and humoral immune responses in mice in comparison with the parental MVA-B vector without deletions, and induced the expression of type I IFN and IFN-α/β inducible genes in human macrophages and monocyte-derived dendritic cells (moDCs) (22, 24). Moreover, vaccination with the VACV strain Western Reserve (WR) lacking the *C6L* gene provided better protection against a challenge with a lethal dose of WR, and induced an enhanced immunogenicity (25).

We have previously described a vaccine candidate against HCV based on MVA strain constitutively expressing the nearly full-length HCV genome from genotype 1a (termed MVA-HCV). In vaccinated mice, MVA-HCV induced polyfunctional HCV-specific CD8^+^ T cell immune responses, mainly directed against p7+NS2 and NS3. Moreover, MVA-HCV induced memory T cell responses with an effector memory phenotype (26). With the aim to enhance the immune responses of MVA-HCV and due to the importance of IFN in HCV infection, we reasoned that a vaccine able to induce the activation of an IFN response might surpass the inhibitory effects of the HCV proteins and thus, improved immune responses to HCV. For this goal, we deleted in MVA-HCV the VACV *C6L* gene, coding for an inhibitor of IFN-β, and performed a head-to-head comparison between MVA-HCV and MVA-HCV ΔC6L analyzing the expression of HCV proteins and evaluating by real-time polymerase chain reaction (PCR) and microarrays the profile of host gene expression induced after infection of human moDCs or macrophages. Furthermore, we have analyzed the innate immune responses *in vivo* in mice inoculated with MVA-HCV and MVA-HCV ΔC6L, together with the adaptive and memory HCV-specific T cell and humoral immune responses. Our findings revealed that both MVA-HCV vectors are capable of activating HCV-specific CD8^+^ T cell and humoral immune responses in spite of the suppressive transcriptional effects mediated by HCV proteins.

## RESULTS

### Generation of MVA-HCV ΔC6L deletion mutant

We have previously described that deletion of the immunomodulatory VACV *C6L* gene, encoding an inhibitor of IFN-β (21, 23), in the vector backbone of an MVA-based HIV/AIDS vaccine candidate induced the production of IFN-β and type I IFN inducible genes in infected innate immune cells and elicited an enhancement in the HIV-specific immunogenicity in immunized mice (22, 24). Thus, to examine whether VACV *C6L* gene could also influence the type I IFN production and the immunogenicity profile of HCV antigens delivered from a poxvirus vector, we deleted *C6L* from the vector backbone of the HCV vaccine candidate MVA-HCV (expressing the nearly full-length genome from HCV genotype 1a) (26), generating the recombinant MVA-HCV ΔC6L deletion mutant (Fig. 1A), as described in materials and methods.

**Figure 1.**
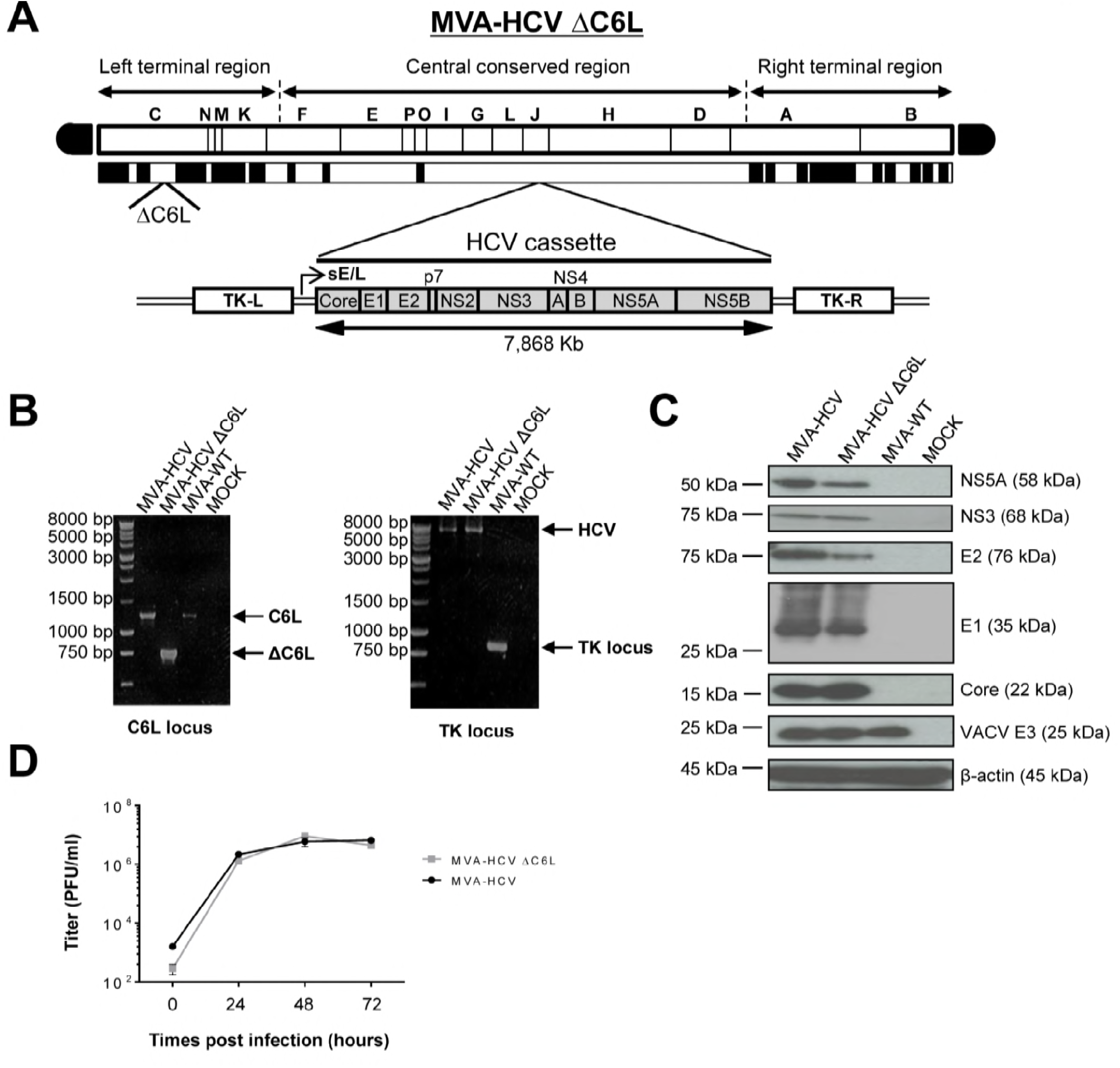
Generation and *in vitro* characterization of MVA-HCV ΔC6L. (A) Scheme of the MVA-HCV ΔC6L deletion mutant genome map. The different regions of the MVA vector are shown in capital letters. Below the map, the deleted or fragmented VACV genes are depicted as black boxes, with the deletion of *C6L* being indicated. The organization of the HCV genome (genotype 1a, strain H77), driven by the sE/L VACV promoter and inserted within the VACV TK viral locus (J2R), is indicated. TK-L, TK left; TK-R, TK right. (B) PCR analysis of the VACV *C6L (*left panel) and TK (right panel) loci. DNA products are indicated by an arrow on the right. (C) Expression of HCV proteins in DF-1 cells infected with MVA-HCV or MVA-HCV ΔC6L at 24 hpi. (D) Viral growth kinetics in DF-1 cells infected with MVA-HCV or MVA-HCV ΔC6L. At different times postinfection, cells were harvested and virus titers in cell lysates were determined by plaque immunostaining assay with anti-VACV antibodies. The mean and standard deviation (SD) from two independent experiments is shown.

The correct generation (scheme in Fig. 1A) and purity of MVA-HCV ΔC6L was confirmed by PCR, as it is shown in Fig. 1B. Viral DNA was purified from DF-1 cells infected with MVA-HCV ΔC6L (or parental MVA-HCV and wild-type attenuated MVA [MVA-WT], used as controls) and amplified by PCR using a set of primers annealing in the *C6L* flanking regions, to confirm the presence of the deletion in the VACV *C6L* gene. The results showed that MVA-HCV ΔC6L has correctly deleted the VACV *C6L* gene (Fig. 1B, left). Moreover, we confirmed the correct insertion of the HCV genes by PCR analysis using a set of primers annealing in the VACV thymidine kinase (TK) flanking sequences, with MVA-HCV and MVA-HCV ΔC6L having a full-length size of about 8 Kb (Fig. 1B, right), that was also confirmed by DNA sequencing of the inserted HCV genome (data not shown). Thus, MVA-HCV ΔC6L vector was successfully generated, with deletion of the VACV *C6L* gene and insertion of HCV genes in the MVA TK locus, and without the presence of wild-type virus.

### Expression of HCV proteins by MVA-HCV ΔC6L

To verify that MVA-HCV ΔC6L constitutively expresses and correctly processes the HCV polyprotein, we performed a Western blot analysis of DF-1 cells infected with MVA-HCV ΔC6L, MVA-HCV or MVA-WT, using specific antibodies that recognized the HCV Core, E1, E2, NS3 and NS5A proteins. As shown in Fig. 1C, the HCV open reading frame was efficiently transcribed and translated during MVA-HCV ΔC6L infection, as did with the parental virus MVA-HCV; producing a viral polyprotein that is correctly processed into mature structural (Core, E1 and E2) and nonstructural (p7-NS2-NS3-NS4-NS5) HCV proteins.

### VACV *C6L* gene is non-essential for MVA-HCV growth

We have previously shown that MVA-HCV and MVA-WT grow similarly in cell culture (26). Here, to further determine whether deletion of VACV *C6L* gene affects virus replication in cell culture, we next compared the virus growth of MVA-HCV ΔC6L and MVA-HCV in DF-1 cells. The results showed that the kinetics of viral growth are similar between parental MVA-HCV and MVA-HCV ΔC6L deletion mutant (Fig. 1D), indicating that the *C6L* gene deletion does not impair virus replication under permissive conditions, and *C6L* is not required for MVA-HCV replication. Furthermore, the mere isolation of MVA-HCV ΔC6L deletion mutant demonstrates that the VACV C6 protein is not essential for MVA-HCV replication.

### RT-PCR showed that MVA-HCV and MVA-HCV ΔC6L downregulate expression of genes involved in innate immunity

We have previously reported that infection of cells with MVA-HCV reduced the expression of several genes involved in innate immunity compared to MVA-WT (26). Thus, to examine whether C6 VACV type I IFN inhibitor protein is able to impair the response of innate immune cells to MVA-HCV promoting an enhancement in the production of type I IFN, we analyzed the expression of type I IFN (IFN-β), IFN-β-induced genes (IFIT1 and IFIT2) and the proinflammatory cytokine TNF-α, by real-time PCR in human THP-1 cells (Fig. 2A) or moDCs (Fig. 2B) mock infected or infected for 6 h with MVA-WT, MVA-HCV and MVA-HCV ΔC6L. The results showed that, compared to MVA-WT, the recombinant viruses MVA-HCV and MVA-HCV ΔC6L similarly down-regulated the expression of IFN-β, IFIT1, IFIT2 and TNF-α in both human cell types without significant differences between both vectors, while mRNA levels were upregulated by MVA-WT.

**Figure 2.**
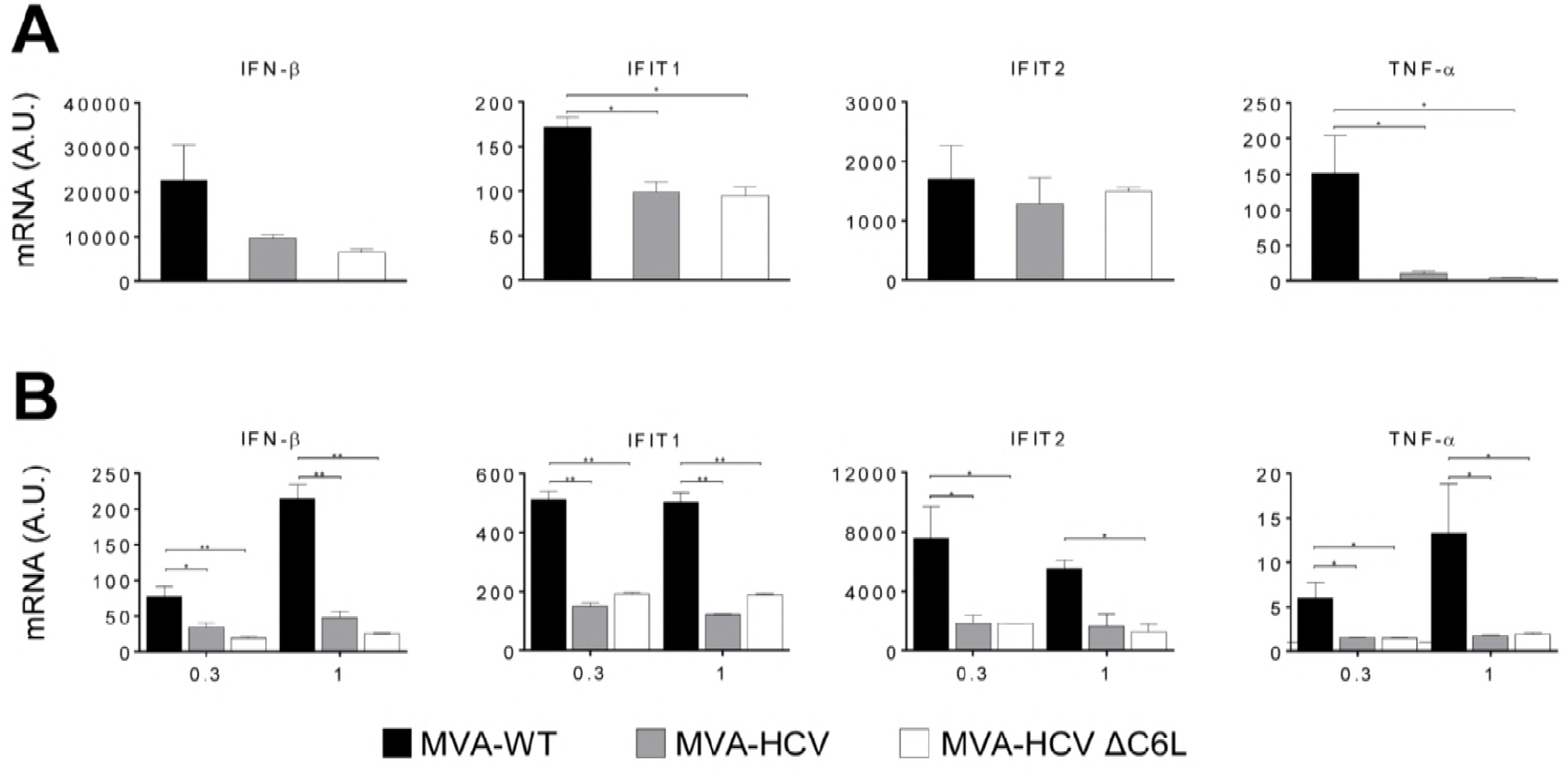
Innate immune responses induced by MVA-HCV and MVA-HCV ΔC6L in human macrophages and moDCs. Human THP-1 macrophages (A) and moDCs (B) were mock infected or infected with MVA-WT, MVA-HCV, or MVA-HCV ΔC6L (5 PFU/cell in panel A and 0.3 or 1 PFU/cell in panel B). At 6 hpi, RNA was extracted and IFN-β, IFIT1, IFIT2, TNF-α and HPRT mRNA levels were analyzed by RT-PCR. Results are expressed as the ratio of the gene to HPRT mRNA levels. A.U., arbitrary units. P values indicate significant response differences when comparing MVA-HCV and MVA-HCV ΔC6L with MVA-WT (* p<0.05, ** p<0.005, *** p<0.001). Data are means ± SD of duplicate samples from one experiment and are representative of two independent experiments.

### Microarray analysis revealed a severe reduction of host gene expression by MVA-HCV and MVA-HCV ΔC6L

To further define whether the two MVA-HCV vectors impact similarly or differently on host gene expression, we performed a microarray analysis to study the differentially expressed host genes in moDCs obtained from three human donor samples and mock infected or infected with MVA-HCV, MVA-HCV ΔC6L and MVA-WT at 1 PFU/cell after 6 h of infection. Under stringent conditions for the microarray analysis and compared to mock infected cells, the results showed that 17 host genes were significantly upregulated in moDCs infected with MVA-WT (Fig. 3A, black bars). In the case of cells infected with MVA-HCV or MVA-HCV ΔC6L host gene expression was still upregulated but reduced several-fold in most of the genes when compared to MVA-WT (Fig. 3A, grey and white bars); and only one gene (TFEB) was downregulated by the three vectors (Fig. 3A). Genes significantly downregulated in MVA-HCV and MVA-HCV ΔC6L compared to MVA-WT include IFN-related genes (OASL, ZC3HAV1, IFN-B1, IFIT1, IFIT2 and IFIT3), genes involved in apoptosis (PMAIP1) and histones (HIST1H4D, HIST1H4F, HIST1H4K, HIST2H4B, HIST1H4H and HIST1H4J). Additionally, in Table 1 we show the host genes that were found differentially expressed in MVA-HCV ΔC6L compared to MVA-HCV, with most of the genes downregulated by MVA-HCV ΔC6L in comparison to MVA-HCV.

**Table 1:** List of genes differentially regulated in human moDCs infected with MVA-HCV ΔC6L in comparison with MVA-HCV. ^a^ The gene name and the probe ID are indicated. ^b^ Differentially expressed genes were evaluated by the non-parametric algorithm ‘Rank Products’ available as “RankProd” package at Bioconductor, and the genes were filtered using a P value Limma <0.05 and a fold change ≥ 2 or ≤ −2.

| Gene Name (probe ID)<sup>a</sup> | Accession number | Fold Change<sup>b</sup> |
| --- | --- | --- |
| Transmembrane protein 11 (TMEM11) | NM_003876 | +2.05 |
| KIAA0586 (KIAA0586) | NM_014749 | +2.00 |
| CD28 molecule (CD28) | NM_006139 | -2.00 |
| FLJ30313 (lnc-GATA5-2) | lnc-GATA5-2:2 | -2.01 |
| Aquaporin 1 (AQP1) | NM_198098 | -2.03 |
| Gessler Wilms tumor Homo sapiens (SMCR2) | AI821758 | -2.05 |
| Oogenesis homeobox (NOBOX) | NM_001080413 | -2.05 |
| RAS oncogene family (RAB6A) | NM_002869 | -2.08 |
| Ribosomal protein S28 (RPS28) | NM_001031 | -2.08 |
| Major histocompatibility complex, class II, DR beta 5 (HLA-DRB5) | NM_002125 | -2.09 |
| Chimerin 2 (ENST00000409922) | ENST00000409922 | -2.10 |
| Nuclear factor of activated T-cells, cytoplasmic, calcineurin-dependent 1 (NFATC1) | NM_172390 | -2.13 |
| RAS guanyl releasing protein 2 (RASGRP2) | ENST00000377494 | -2.14 |
| Glycosyltransferase 3 family (UGT3A1) | NM_152404 | -2.16 |
| Myotubularin related protein 3 (MTMR3) | NM_021090 | -2.17 |
| Keratin associated protein 2-1 (KRTAP2-1) | NM_001123387 | -2.18 |
| keratin associated protein 5-4 (KRTAP5-4) | NM_001012709 | -2.19 |
| G protein-coupled receptor 88 (GPR88) | NM_022049 | -2.19 |
| G protein pathway suppressor 2 pseudogene (ENST00000430745) | ENST00000430745 | -2.19 |
| Secretory trafficking family member B (MON1B) | NM_014940 | -2.21 |
| Solute carrier family 16, member 11 (SLC16A11) | NM_153357 | -2.24 |
| Vacuolar protein sorting 18 homolog (VPS18) | NM_020857 | -2.26 |
| Synapse defective 1, Rho GTPase, homolog 2 (SYDE2) | NM_032184 | -2.27 |
| Neuron-derived neurotrophic factor (NDNF) | NM_024574 | -2.30 |
| Homeobox 1 (EMX1) | BC037242 | -2.32 |
| Actin-like protein 3 (THC2499666) | THC2499666 | -2.32 |
| Interleukin 15 receptor (IL15RA) | ENST00000379971 | -2.39 |
| Chemokine (C-C motif) ligand 18 (CCL18) | NM_002988 | -2.40 |
| cDNA clone NT2RI2025693 (ENST00000587777) | ENST00000587777 | -2.42 |
| Cellular repressor of E1A-stimulated genes 1 (CREG1) | NM_003851 | -2.46 |
| NYN domain and retroviral integrase containing (NYNRIN) | NM_025081 | -2.49 |
| Uncoupling protein 3 (UCP3) | NM_022803 | -2.53 |
| Phospholipase C (PLCH2) | NM_014638 | -2.57 |
| Long intergenic non-protein coding RNA 687 (LINC00687) | NR_110635 | -3.05 |
| Synuclein, beta (SNCB) | NM_001001502 | -3.05 |
| Arylformamidase (AFMID) | NM_001145526 | -3.20 |

**Figure 3.**
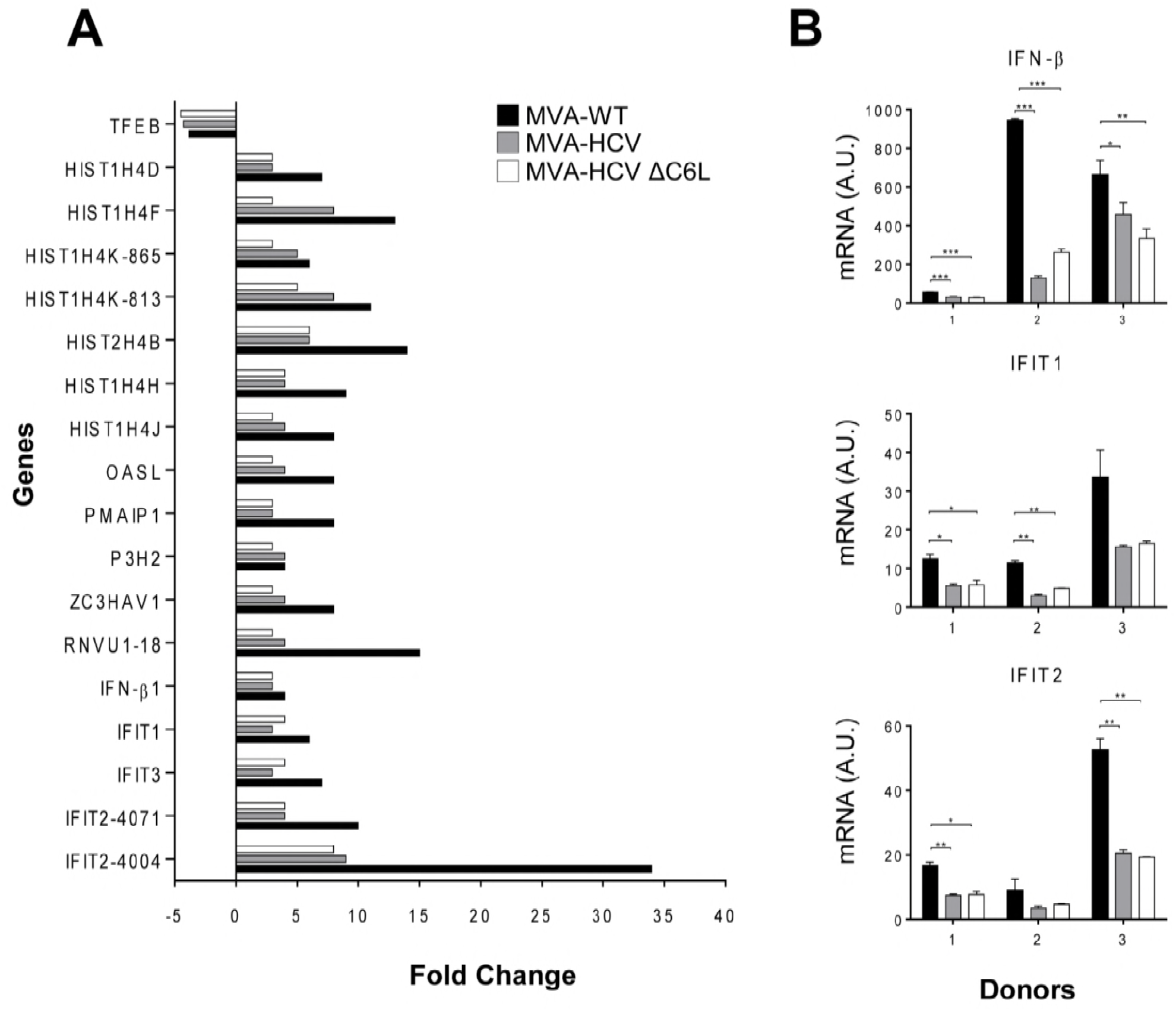
Microarray analysis of HCV antigen-regulated host genes in human moDCs infected with MVA-HCV and MVA-HCV ΔC6L. (A) Clustering classification of host genes differentially regulated in MVA-WT-, MVA-HCV- and MVA-HCV ΔC6L-infected human moDCs versus mock infected cells (FDR Prod p value <0.05, fold change ≥ 2 or ≤ - 2). MoDCs were infected for 6 h and the expression levels of different host genes were analyzed by using a human oligo microarray 4×44K, as described in Material and Methods. Differentially expressed genes were evaluated by the non-parametric algorithm ‘Rank Products’ available as “RankProd” package at Bioconductor. Results are the mean obtained from three different healthy blood donors. (B) Validation of microarray data by real-time RT-PCR to determine the mRNA levels of IFN-β, IFIT1, and IFIT2 genes. RNA isolated from the three healthy blood donors used for microarray analysis was used. A.U., arbitrary units. Data are means ± SD of duplicate samples. P values indicate significant response differences comparing MVA-HCV and MVA-HCV ΔC6L with MVA-WT (* p<0.05).

To validate the microarray results, we performed real-time PCR of the mRNA samples using specific primers for three representative host genes (IFN-β, IFIT1 and IFIT 2) (Fig. 3B). The results obtained were in accordance with the microarray data, showing a downregulation of these genes in MVA-HCV and MVA-HCV ΔC6L, compared to MVA-WT in the three human donor samples.

These results established that in moDCs infected with MVA-HCV and MVA-HCV ΔC6L there is a reduced host gene expression compared to parental MVA-WT, with minor differences in extent between the two recombinant viruses.

### MVA-HCV and MVA-HCV ΔC6L triggered differential cell recruitment than MVA-WT in the peritoneal cavity of infected mice

Due to the impact of MVA-HCV and MVA-HCV ΔC6L on host gene expression, we next determined the pattern of immune cells that migrate *in vivo* to the peritoneal cavity after vector’s exposure. Hence, we inoculated intraperitoneally (i.p.) the different viruses in five C57BL/6 mice per group (MVA-WT, MVA-HCV, MVA-HCV ΔC6L and PBS) as described in Materials and Methods, and analyzed by flow cytometry the absolute numbers of several immune cell populations present in the peritoneal cavity at 6, 24 and 48 h post inoculation (Fig. 4). The results showed a large cell recruitment at this site for the total number of immune cells (around 10^7^ cells) following vector inoculation. However in infections with MVA-HCV and MVA-HCV ΔC6L there was lower recruitment in the total number of specific immune cell types (dendritic cells [DCs], natural killer [NK] cells, NK T [NKT] cells, and CD4^+^ and CD8^+^ T cells) than for MVA-WT at 24 h post inoculation (Fig. 4 and Table 2). At 24 h after viral infection most of the macrophages left the peritoneal cavity, while DCs, NKs, NKT and neutrophils (total, α and β) were highly recruited at 24 h and 48 h in MVA-WT, MVA-HCV and MVA-HCV ΔC6L in comparison with PBS. In absolute numbers, most of the neutrophils recruited with the MVA vectors were neutrophils α. There were no significant differences between MVA-HCV and MVA-HCV ΔC6L at any time point, with only a trend to higher number of recruited NK, NKT cells and neutrophils (total, α and β) in the case of MVA-HCV ΔC6L at 48 h. This tendency is statistically significant in the number of neutrophils recruited (total, α and β) in the comparison of MVA-HCV ΔC6L with MVA-WT (Table 2).

**Table 2:**
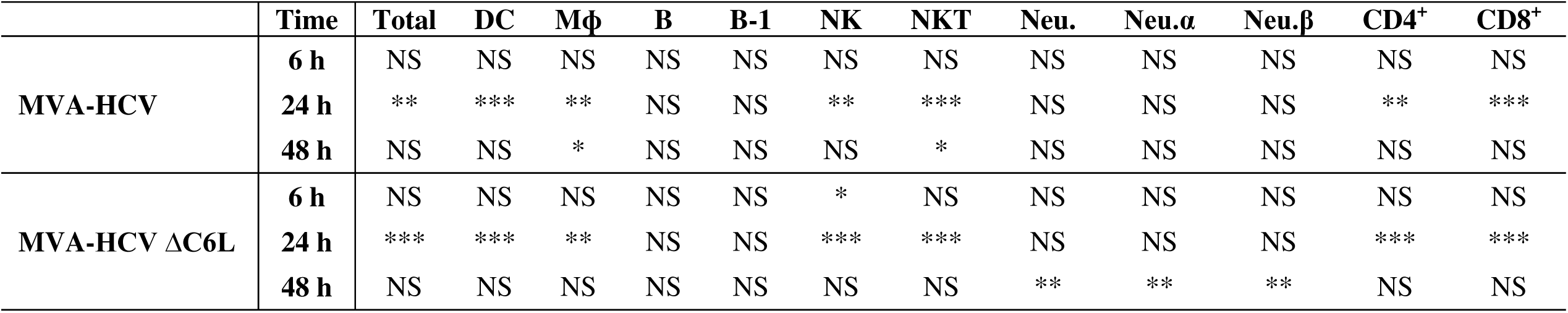
Statistical significances of the comparison between MVA-HCV and MVA-HCV ΔC6L versus MVA-WT in the recruitment of innate immune cells in the peritoneal cavity of infected mice. Results are represented in Figure 4. NS (no significant) =P > 0.05; * =P ≤ 0.05; ** = P ≤ 0.01; *** = P ≤ 0.001. Total (total number of cells), DC (Dendritic Cells), Mϕ (Macrophages), B (B cells), B-1 (B-1 cells), NK (Natural Killer cells), NKT (Natural Killer T cells), Neu. (Neutrophils), CD4^+^ (CD4^+^ T cells), CD8^+^ (CD8^+^ T cells).

**Figure 4.**
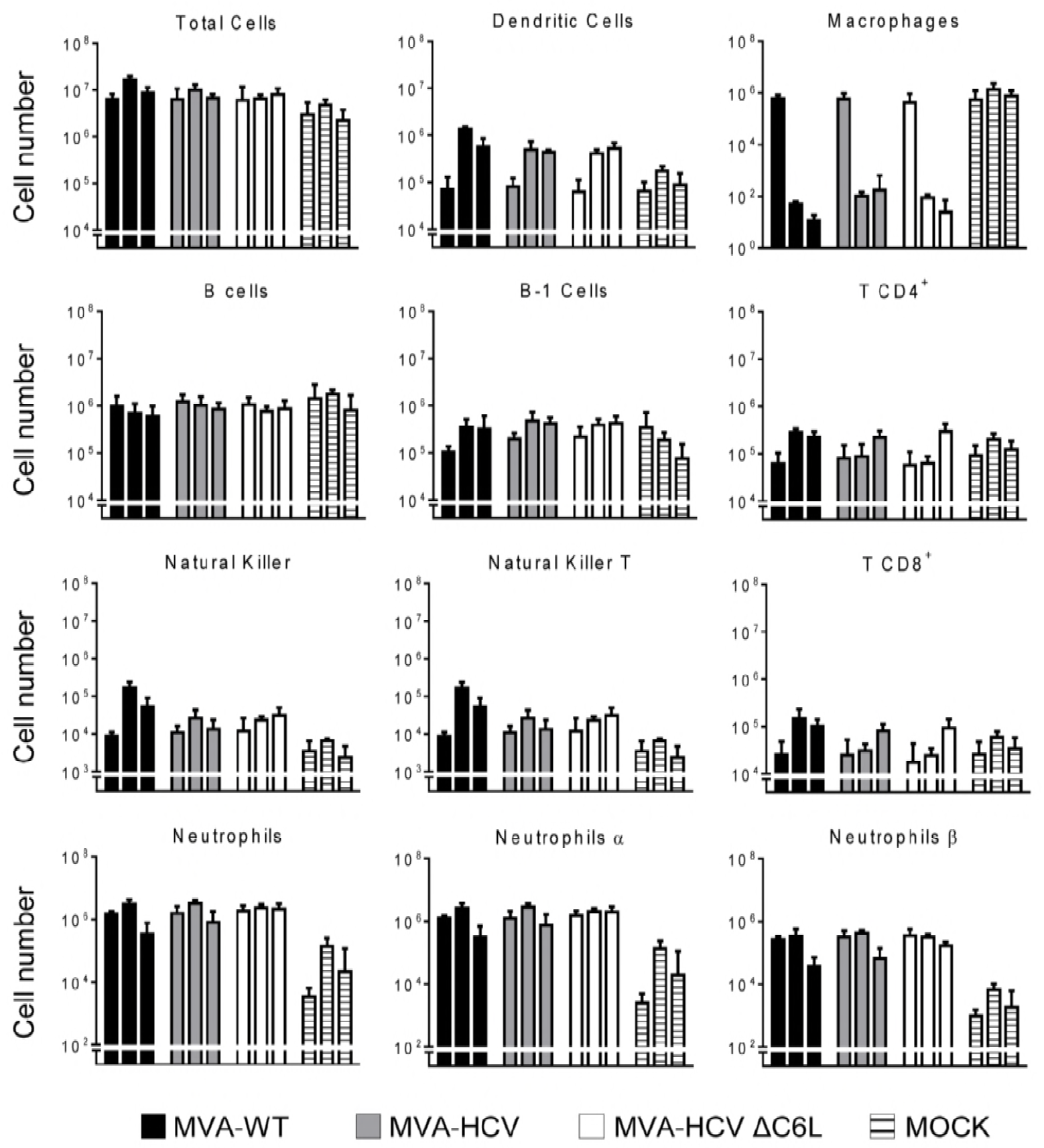
Recruitment of innate immune cells in the peritoneal cavity of infected mice. Absolute numbers of innate immune cell populations obtained from the peritoneal cavity of C57BL/6 mice infected by the i.p. route with MVA-WT, MVA-HCV, MVA-HCV ΔC6L or PBS. Peritoneal exudate cells were collected at 6, 24 and 48 h post inoculation (first, second and third bar in each group, respectively) from each individual mouse (n=5 per group), stained for different surface markers and absolute numbers analyzed by flow cytometry. Graphs show the logarithmic mean ± SD. Statistical significance between groups was calculated with a t-student test (unpaired, non-parametric, two tails). P values are indicated in Table 2.

These results showed the impact of the MVA vectors on immune cell recruitment in the peritoneal cavity of mice during virus infection, revealing statistical differences in some innate cell recruitment between them.

### MVA-HCV and MVA-HCV ΔC6L induced high, broad and polyfunctional HCV-specific T cell adaptive immune responses in immunized mice

To define whether the deletion of C6 VACV IFN-β inhibitor from MVA-HCV could have an *in vivo* impact on the adaptive immune response against HCV antigens, we analyzed the HCV-specific T cell response elicited by MVA-HCV and MVA-HCV ΔC6L in mice immunized following two inoculations with each viral vector (homologous prime-boost immunization protocol). Thus, eight C57BL/6 mice of each group (MVA-WT, MVA-HCV and MVA-HCV ΔC6L) were immunized as described in Materials and Methods and half of them (n=4) were sacrificed at day 10 post-boost to measure by intracellular cytokine staining (ICS) the HCV-specific adaptive T cell immune responses. Pools of splenocytes from each group were stimulated *ex vivo* with a panel of HCV peptides covering the entire sequence of HCV H77 strain (genotype 1a) and after 6 h of stimulation cells were stained with specific antibodies to identify T cell populations (CD4 and CD8) and responding cells (CD107a on the surface of activated T cells as an indirect marker of cytotoxicity, and production of IFN-γ, TNF-α, and IL-2 cytokines).

The magnitude of the total HCV-specific CD8^+^ T cell adaptive immune responses (determined as the sum of cells producing IFN-γ, TNF-α, and/or IL-2, as well as the expression of CD107a degranulation marker) obtained for all the HCV peptide pools (Core + E1 + E2 + p7+NS2 + NS3 + NS4 + NS5) was high and similar in mice immunized with MVA-HCV and MVA-HCV ΔC6L. The immune response was largely mediated by HCV-specific CD8^+^ T cells, while levels of HCV-specific CD4^+^ T cells were very low (Fig. 5A; and data not shown).

**Figure 5.**
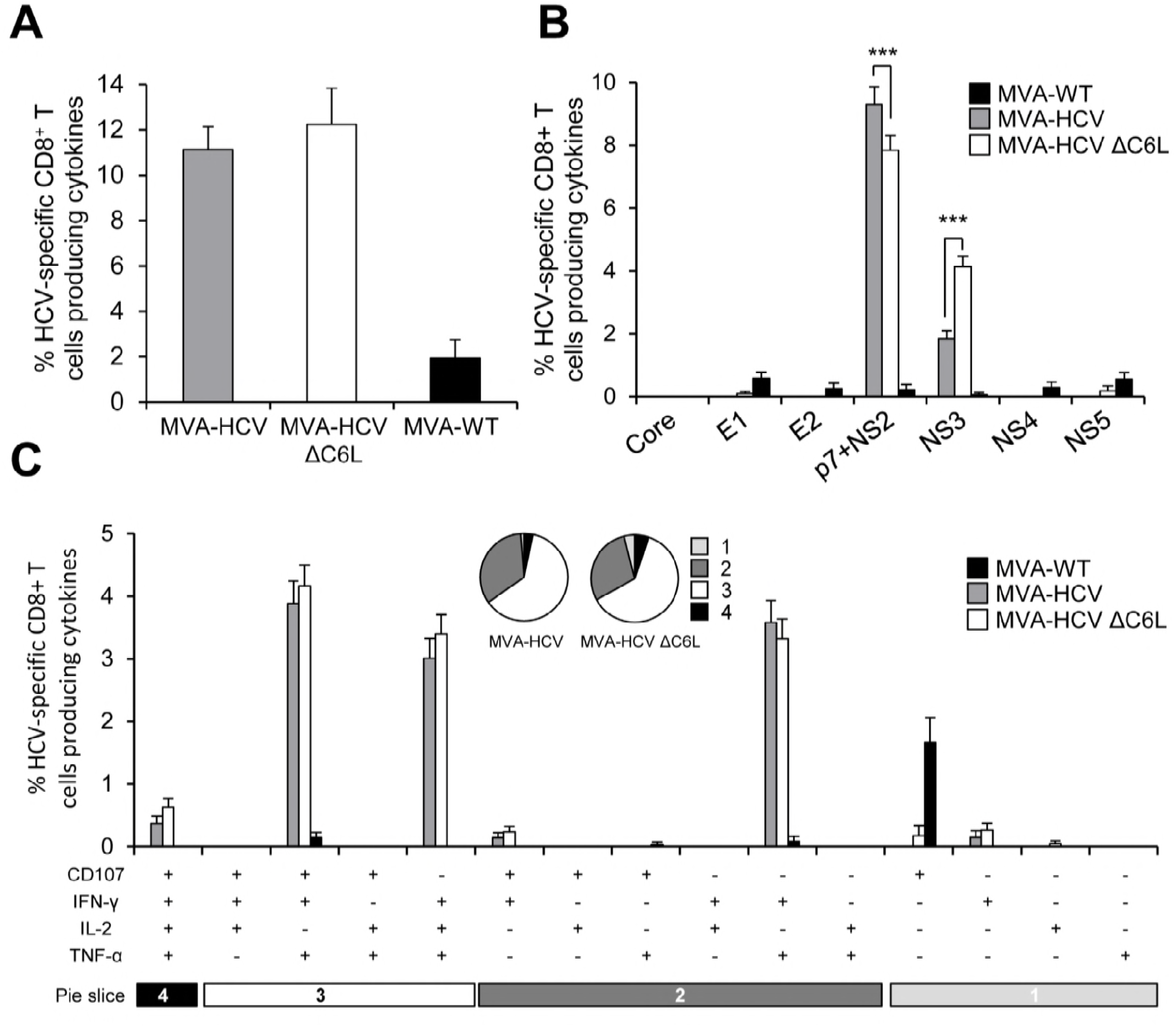
Magnitude, breath and polyfunctionality of HCV-specific CD8^+^ T cell adaptive immune responses. Splenocytes were obtained from mice (n=4 per group) immunized with two doses of MVA-WT, MVA-HCV or MVA-HCV ΔC6L 10 days after the last immunization. Then, HCV-specific CD8^+^ T cell adaptive immune responses elicited were measured by ICS after stimulation of splenocytes with different HCV peptide pools. Values from unstimulated controls were subtracted in all cases. P values indicate significantly response differences when comparing MVA-HCV with MVA-HCV ΔC6L (*** = P ≤ 0.001). Data is from one experiment representative of four independent experiments. (A) Percentage of total HCV-specific CD8^+^ T cell adaptive immune responses directed against HCV antigens. The values represent the sum of the percentages of CD8^+^ T cells producing CD107a and/or IFN-γ and/or TNF-α and/or IL-2 against Core, E1, E2, p7+NS2, NS3, NS4 and NS5 peptide pools. (B) Percentages of Core, E1, E2, p7+NS2, NS3, NS4 or NS5 HCV-specific CD8^+^ T cells. Frequencies represent the sum of the percentages of CD8^+^ T cells producing CD107a and/or IFN-γ and/or TNF-α and/or IL-2 against each HCV peptide pool. (C) Polyfunctionality profile of total HCV-specific CD8^+^ T cell adaptive immune responses directed against HCV antigens. All possible combinations of the responses are shown on the X axis (15 different T cell populations), while the percentages of CD8^+^ T cells producing CD107a and/or IFN-γ and/or TNF-α and/or IL-2 against HCV peptide pools are shown on the Y axis. Responses are grouped and color coded on the basis of the number of functions (one, two, three, or four). Each slice in the pie charts corresponds to the proportion of the total HCV-specific CD8^+^ T cells exhibiting one, two, three, or four functions (CD107a and/or IFN-γ and/or TNF-α and/or IL-2) within the total HCV-specific CD8^+^ T cells.

The profile of HCV-specific CD8^+^ T cell adaptive immune responses elicited by MVA-HCV and MVA-HCV ΔC6L immunization groups showed that the response was directed preferentially against p7+NS2 and to a lesser extent toward NS3 in both immunization groups (Fig. 5B). Interestingly, MVA-HCV induced significantly higher p7+NS2-specific CD8^+^ T cell immune responses than MVA-HCV ΔC6L, while MVA-HCV ΔC6L elicited higher NS3-specific CD8^+^ T cell immune responses than MVA-HCV (Fig. 5B).

The quality of the HCV-specific CD8^+^ T cell immune responses was analyzed measuring the pattern of cytokine production (IFN-γ, TNF-α, and/or IL-2) and/or its cytotoxic potential (CD107a degranulation). As shown in Fig. 5C (pie charts), HCV-specific CD8^+^ T cell immune responses were similar and highly polyfunctional in animals immunized with MVA-HCV and MVA-HCV ΔC6L, with around 60% and 5% of the CD8^+^ T cells having three and four functions, respectively. HCV-specific CD8^+^ T cells producing CD107a + IFN-γ + TNF-α, IFN-γ + TNF-α + IL-2 or IFN-γ + TNF-α were the most abundant cell populations induced by MVA-HCV and MVA-HCV ΔC6L, which were of a similar magnitude between both immunization groups (Fig. 5C, bars).

These results indicate that there are no significant differences in the magnitude and quality of the total HCV-specific CD8^+^ T cell adaptive immune responses induced by MVA-HCV and MVA-HCV ΔC6L. However, the pattern of response was different; with MVA-HCV inducing higher p7+NS2-specific CD8^+^ T cell responses than MVA-HCV ΔC6L and MVA-HCV ΔC6L eliciting higher NS3-specific CD8^+^ T cell immune responses than MVA-HCV.

### MVA-HCV and MVA-HCV ΔC6L induced T cell memory immune responses

It has been reported that memory CD8^+^ T cells are required for protection from persistent HCV (27). Thus, to study the HCV-specific T cell memory immune responses induced by MVA-HCV and MVA-HCV ΔC6L, four mice of each immunization group were sacrificed 53 days after the second immunization and pools of splenocytes from each group were stimulated *ex vivo* with a panel of HCV peptides, similarly to the adaptive immune response protocol described above and in Materials and Methods.

The magnitude of the total HCV-specific CD8^+^ T cell memory immune responses were high and similar in mice immunized with MVA-HCV and MVA-HCV ΔC6L; with the HCV-specific T cell memory immune responses largely mediated by CD8^+^ T cells, with very low HCV-specific CD4^.+^ T cells (Fig. 6A; and data not shown). Moreover, as in adaptive phase, the pattern of HCV-specific CD8^+^ T cell memory immune responses induced by MVA-HCV and MVA-HCV ΔC6L showed that in both groups the response was mainly directed against p7+NS2 and then towards NS3 (Fig. 6B). MVA-HCV elicited significantly higher p7+NS2-specific CD8^+^ T cell immune responses than MVA-HCV ΔC6L, and MVA-HCV ΔC6L elicited higher NS3-specific CD8^+^ T cell immune responses than MVA-HCV (Fig. 6B).

**Figure 6.**
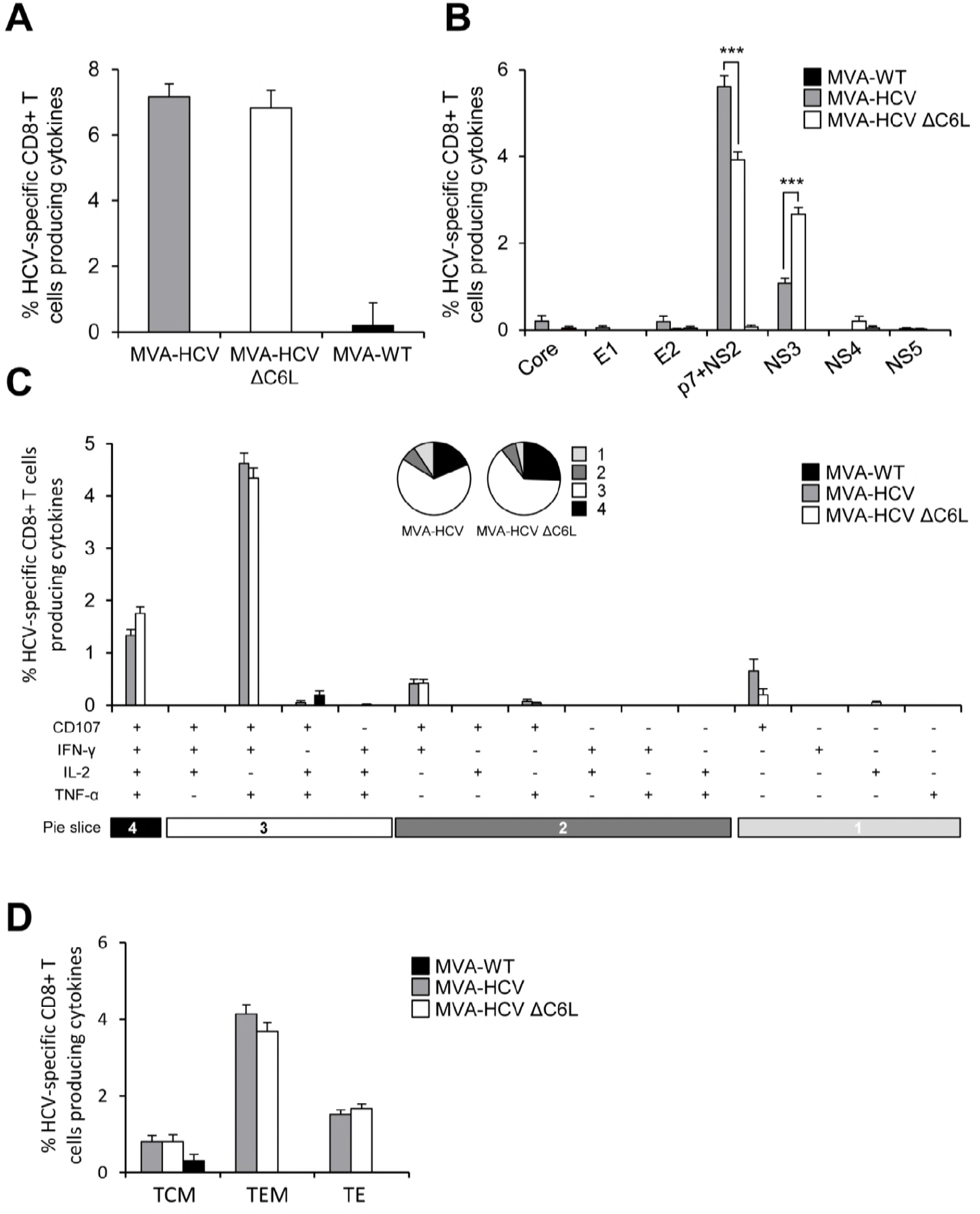
Magnitude, breath, polyfunctionality and phenotype of HCV-specific CD8^+^ T cell memory immune responses. Splenocytes were obtained from mice (n=4 per group) immunized with two doses of MVA-WT, MVA-HCV or MVA-HCV ΔC6L 53 days after the last immunization. Then, HCV-specific CD8^+^ T cell memory immune responses elicited were measured by ICS after stimulation of splenocytes with different HCV peptide pools. Values from unstimulated controls were subtracted in all cases. P values indicate significantly response differences when comparing MVA-HCV with MVA-HCV ΔC6L (*** = P ≤ 0.001). Data is from one experiment representative of four independent experiments. (A) Percentage of total HCV-specific CD8^+^ T cell memory immune responses directed against HCV antigens. The values represent the sum of the percentages of CD8^+^ T cells producing CD107a and/or IFN-γ and/or TNF-α and/or IL-2 against Core, E1, E2, p7+NS2, NS3, NS4 and NS5 peptide pools. (B) Percentages of Core, E1, E2, p7+NS2, NS3, NS4 or NS5 HCV-specific CD8^+^ T cells. Frequencies represent the sum of the percentages of CD8^+^ T cells producing CD107a and/or IFN-γ and/or TNF-α and/or IL-2 against each HCV peptide pool. (C) Polyfunctionality profile of total HCV-specific CD8^+^ T cell memory immune responses directed against HCV antigens. All possible combinations of the responses are shown on the X axis (15 different T cell populations), while the percentages of CD8^+^ T cells producing CD107a and/or IFN-γ and/or TNF-α and/or IL-2 against HCV peptide pools are shown on the Y axis. Responses are grouped and color coded on the basis of the number of functions (one, two, three, or four). Each slice in the pie charts corresponds to the proportion of the total HCV-specific CD8^+^ T cells exhibiting one, two, three, or four functions (CD107a and/or IFN-γ and/or TNF-α and/or IL-2) within the total HCV-specific CD8^+^ T cells. (D) Phenotypic profile of total HCV-specific CD8^+^ T cell memory immune responses directed against HCV antigens. Frequencies represent percentages of TCM, TEM, and TE HCV-specific CD8^+^ T cells producing CD107a and/or IFN-γ and/or TNF-α and/or IL-2 against HCV peptide pools. TCM, TEM, and TE phenotypes are determine on the basis of CD127 and CD62L expression: TCM (CD127^+^, CD62L^+^), TEM (CD127^+^, CD62L^−^), and TE (CD127^−^, CD62L^−^).

The quality of the HCV-specific CD8^+^ T cell memory immune responses showed that MVA-HCV and MVA-HCV ΔC6L triggered highly polyfunctional HCV-specific CD8^+^ T cells, with around 65% and 23% of the CD8^+^ T cells having three and four functions, respectively (Fig. 6C, pie charts). HCV-specific CD8^+^ T cells producing CD107a + IFN-γ + TNF-α + IL-2 and CD107a + IFN-γ + TNF-α were the most abundant cells elicited by MVA-HCV and MVA-HCV ΔC6L (Fig. 6C, bars).

Additionally, we examined the phenotype of the total HCV-specific CD8^+^ T memory cells by measuring the expression of the CD127 and CD62L surface markers, which allow the definition of the different memory subpopulations: T central memory (TCM, CD127^+^/CD62L^+^), T effector memory (TEM, CD127^+^/CD62L^−^), and T effector (TE, CD127^−^/CD62L^−^) cells (Fig. 6D). The results showed that immunization with MVA-HCV and MVA-HCV ΔC6L elicited a high percentage of HCV-specific CD8^+^ T memory cells, which were mainly of the TEM phenotype, and of similar magnitude in both immunization groups.

These results indicate that there are no significant differences in the magnitude and quality of the total HCV-specific CD8^.+^ T cell memory immune responses induced by MVA-HCV and MVA-HCV ΔC6L. However, the pattern of response was again different; with MVA-HCV inducing higher p7+NS2-specific CD8^+^ T cell responses than MVA-HCV ΔC6L and MVA-HCV ΔC6L eliciting higher NS3-specific CD8^+^ T cell immune responses than MVA-HCV.

### MVA-HCV and MVA-HCV ΔC6L induced antibodies against HCV E2 protein

Although the role of the humoral immune response in controlling HCV infection has been controversial for many years, it has been described that passive transfer of anti-E2 antibodies protects against infection in chimpanzees and chimeric mice (28–30). Thus, we analyzed the humoral immune responses induced after immunization with MVA-HCV and MVA-HCV ΔC6L, quantifying by enzyme-linked immunosorbent assay (ELISA) the total IgG levels of antibodies against HCV E2 protein (strain H77) in sera obtained from individual mice 10 days post-boost (Fig. 7). The results showed that both immunization groups were able to elicit antibodies against E2 in the adaptive phase.

**Figure 7.**
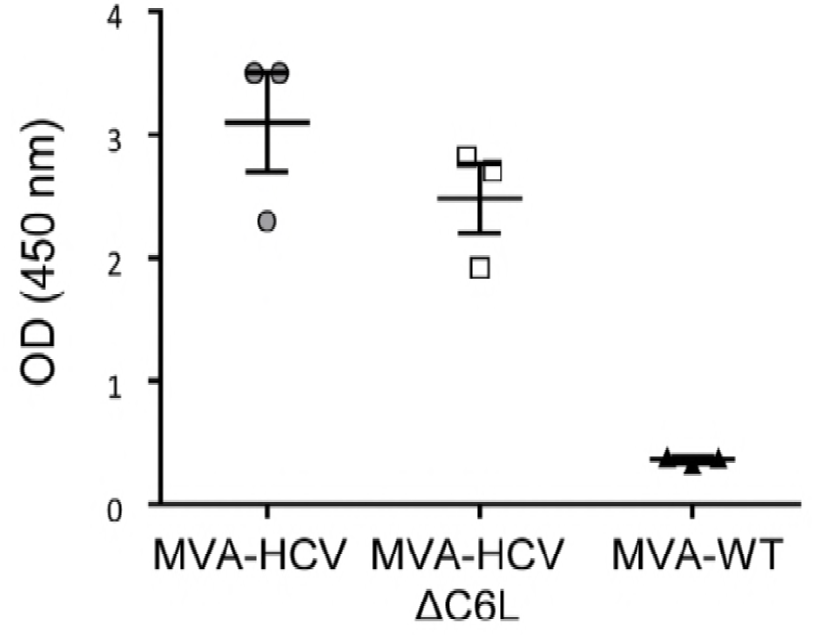
Humoral immune responses elicited by MVA-HCV and MVA-HCV ΔC6L against HCV E2 protein. Levels of E2-specific total IgG binding antibodies were measured by ELISA in serum from individually mice immunized with two doses of MVA-WT, MVA-HCV or MVA-HCV ΔC6L at day 10 after the last immunization. Absorbance values (measured at 450 nm) correspond to 1/100 dilution of individual serum, and each mouse is represented by a dot. The mean ± SD are indicated.

## DISCUSSION

In spite of the great advances in the development of effective anti-HCV drugs, and due to a worldwide large population of infected persons, undoubtedly there is the need to develop an HCV vaccine. An ideal vaccine against HCV should be able to cure the disease during primary infection and prevent chronicity, since HCV-associate diseases are mostly manifested during chronic infection where treatment is less successful and liver is being severely damaged (31). Spontaneous cure of infection happens in 30% of infected people, resulting in a complete viral clearance and higher chances of resolving a reinfection, staying away from chronicity. Therefore, it is clear that some components of the immune system are able to skew the outcome of HCV infection towards resolution after the acute phase, making this the main goal of a successful HCV vaccine (11).

Over the last few years several HCV vaccines candidates aimed to induce strong humoral responses and/or T cell responses have been tested in preclinical studies, but only a few of them reached clinical trials (31). Several viral vector-based vaccines have been tested in humans, including adenovirus vectors expressing the nonstructural HCV proteins NS3, NS4 and NS5mut (genetically inactivated polymerase gene) (Ad6NSmut and AdCh3NSmut) in Phase I clinical trials (study HCV001 NCT01070407 in healthy volunteers and study HCV002 NCT01094873 in chronically infected patients), showing that both vectors are safe and immunogenic (32). Furthermore, an MVA expressing the same antigens (MVA-NSmut) was also assessed alone or in combination with Ad6NSmut (NCT01701336 in chronically infected patients) or AdCh3NSmut (NCT01296451 in healthy volunteers and HCV infected patients) in Phase I clinical trials (33). Currently, AdCh3NSmut and MVA-NSmut are being tested in a Phase I/II clinical trial (NCT01436357 in uninfected injection drug users, final data collection in July, 2018). Lastly, an MVA vector encoding NS3, NS4 and NS5b (TG4040) was studied in a dose-escalating Phase I clinical trial (NCT00529321 in untreated chronic patients), showing that this vector is also safe and immunogenic. TG4040 was further tested in a Phase II clinical trial as a therapeutic vaccine combined with standard of care treatment (PEG-IFN-α with ribavirin) (NCT01055821 in untreated chronic patients) (34). In clinical and preclinical trials, both adenoviruses and MVA vectors elicited HCV-specific CD8^+^ T cell responses; however, due to the anti-adeno sero-prevalence in human population and the cross-reactive immunity between both vectors (AdCh3NSmut and Ad6NSmut), the immunogenicity was improved when MVA was administered as a boost (33). Moreover, a recent study showed that in immunized mice an MVA vector expressing NS3/4A is superior to adenovirus-5 vector in the induction of CD8^+^ T cell memory responses against HCV (35).

We have previously described an MVA-based vaccine candidate for HCV (termed MVA-HCV) that expresses most of the HCV genome (Core, E1, E2, p7, NS2, NS3, NS4a, NS4b, NS5a and part of NS5b protein) (26). MVA-HCV elicited polyfunctional T cell responses with memory phenotype in wild-type and humanized mice, being the first vaccine candidate encoding all HCV proteins (structural and nonstructural) and therefore covering most of the T and B cell determinants described for HCV. However, efforts to understand the HCV-specific immune responses elicited by MVA-HCV are needed in order to improve this vaccine candidate. Firstly, despite of its attenuated phenotype, MVA still encodes for proteins that can interfere with host immune responses to viral infection (36–38), and it is described that deletion of VACV immunomodulatory genes can enhance its immunogenicity (20). Secondly, IFN-based therapy in combination with ribavirin has been the treatment of first choice despite its well-known side effects, and even in the era of new DAA therapy, treatment with IFN results in improvement of response rates and higher sustained virological response in genotypes 1 (39, 40), 2 and 3 (41, 42). Therefore, IFN release and a strong immune response are critical in the control and resolution of HCV infection providing signals for the efficient priming of the adaptive branch of immune response (19).

Hence, in an effort to define the immunomodulatory role of an IFN response over the MVA-HCV vaccine candidate, we deleted the VACV *C6L* gene (encoding a type I IFN inhibitor protein) from MVA-HCV to generate a novel HCV vaccine candidate termed MVA-HCV ΔC6L. MVA-HCV ΔC6L efficiently produces all HCV antigens (Core, E1, E2, p7, NS2, NS3, NS4a, NS4b, NS5a and a part of NS5b) at the same level as MVA-HCV during the course of virus infection. MVA-HCV ΔC6L replicates similarly to MVA-HCV in permissive culture cells, indicating that deletion of *C6L* had no effect in viral kinetics. Unexpectedly, MVA-HCV ΔC6L did not increase in infected human macrophages or moDCs the expression of type I IFN or IFN-related genes compared to parental MVA-HCV. Although both vectors downregulated host gene expression in comparison with MVA-WT, they still induced immune activation despite of the presence of HCV proteins, which are potent innate response inhibitors, especially the NS3 HCV protein (43, 44).

Microarray analyses in infected human moDCs provided an identification of the host genes triggered in response to the HCV proteins expressed from MVA-HCV and MVA-HCV ΔC6L. Under stringent conditions of microarray analysis, the results showed that MVA-HCV and MVA-HCV ΔC6L differentially downregulate IFN-related genes (OASL, ZC3HAV1, IFN-B1, IFIT1, IFIT2 and IFIT3), genes involved in apoptosis (PMAIP1) and histones (HIST1H4D, HIST1H4F, HIST1H4K, HIST2H4B, HIST1H4H and HIST1H4J) in comparison with MVA-WT. This downregulation of type I IFN and IFN-related genes confirmed the previous RT-PCR results in infected moDCs, and is in accordance with the role exerted by the HCV proteins, specially NS3, inhibiting type I IFN signaling (43, 44). In a different virus-cell system in which the same HCV cassette was expressed from a transcriptional regulated replication competent VACV-based delivery system (VT7-HCV7.9) in HeLa cells, we previously reported the induction of cell apoptosis by HCV proteins mediated by IFN-induced enzymes PKR and RNase-L (45) and that HCV protein expression modulated transcription of genes associated with lipid metabolism, oxidative stress, apoptosis, and cellular proliferation (46). Down-regulation of histone mRNA levels, which are tightly regulated during DNA replication, indicates that cell proliferation and cell cycle progression are likely impaired in cells infected with MVA-HCV and MVA-HCV ΔC6L. In this regard it has been reported that infection of Huh7.5 cells with HCV resulted in inhibition of histone H4 methylation/acetylation and histone H2AX phosphorylation, with a significant changed in expression of genes important for hepatocarcinogenesis, inhibiting DNA damage repair (47).

In cells recruited in the peritoneal cavity of mice inoculated with MVA-HCV, MVA-HCV ΔC6L, MVA-WT or PBS, we also observed an inhibitory effect in the innate immune responses induced by the MVA vectors expressing HCV proteins, that resulted in less recruitment of specific innate immune cell types, such as DCs, NK, NKT, CD4^.+^ and CD8^+^ T cells at 24 h post-inoculation in comparison with MVA-WT. This lower recruitment of innate immune cells is in accordance with the downregulation of type I IFN and IFN-related genes observed in human moDCs infected with MVA-HCV and MVA-HCV ΔC6L.

We have previously shown that deletion of VACV *C6L* gene in the vector backbone of the MVA-B vaccine candidate against HIV/AIDS, expressing HIV-1 Env, Gag, Pol and Nef antigens (MVA-B ΔC6L), significantly upregulates IFN-β and IFN-α/β-inducible genes (22, 24). Here, the same MVA vector but expressing HCV instead of HIV-1 antigens shows opposite results, likely due to the nature of HCV proteins in inhibiting type I IFN responses. This result highlights the influence of expressing different heterologous antigens in the biology of the MVA vector. It has been described that nonstructural proteins from the *Flaviviridae* family interfere with the host immune responses to facilitate viral propagation (48): HCV NS2 inhibits IRF3 phosphorylation and therefore the production of IFN-β, IFNα1, IFNλ1, IFNλ3, and chemokines CCL5 and CXCL19 (49); HCV NS3/4a blocks TLR3 signaling pathway (44), IRF3 phosphorylation (50) and inhibits RLR signaling (43, 44); HCV NS4b avoids STING accumulation suppressing RLR signaling (51); and HCV NS5 impairs TLR-MyD88 signaling (52), blocks IRF7 nuclear translocation (53) and binds to JAK/STAT (54) and PKR (55) inhibiting their signaling pathway. Thus, in the context of the MVA-HCV vaccine candidate, production of all HCV proteins negatively impacts on host gene expression, and deletion of the VACV C6 type I IFN inhibitor is not enough to counteract and reverse the strong IFN inhibition exerted by the HCV proteins; therefore an increase in IFN-β and IFN-α/β-inducible genes is not obtained.

Despite of the inhibition of type I IFN exerted by infection with MVA-HCV and MVA-HCV ΔC6L, these vectors are still able to induce HCV-specific T and B cell responses in immunized mice. Several studies have proven that a potent T cell response is important for virus clearance (56). It has also been observed that a monospecific HCV-specific CD8^+^ T cell response is associated with viral persistence (57), hence a vaccine against HCV should ideally include most of the HCV antigens, including structural and nonstructural proteins, in order to elicit broad T cell response that can target most of HCV epitopes. Furthermore, HCV has a potent error-prone polymerase, leading to viral mutant scape that can be favored by a narrow epitope-targeting immune response. As shown here, MVA-HCV ΔC6L elicited high, broad and polyfunctional HCV-specific CD8^.+^ T cells at a similar level as MVA-HCV, but the specificity of the response against the HCV antigens was different, with MVA-HCV ΔC6L inducing a significantly higher NS3-specific response while MVA-HCV elicited higher p7+NS2-specific response. The impact of these differences needs to be further studied, but it has been proposed that a response against HCV NS3 protein could be more important in protection due to the role of this protein in HCV infection, with several viral epitopes triggering CD8^+^ T cell responses (58, 59). Moreover, DCs pulsed with HCV NS3 protein induced immune responses and protection from infection in mice inoculated with recombinant VACV expressing NS3 (60), and NS3-specific CD8^+^ T cell responses were observed in a chimpanzee clearing HCV infection (61), reinforcing the important role of NS3-specific responses in HCV. Furthermore, all clinical trials have included the NS3 protein in viral vectors, as mentioned above. Interestingly, both MVA-HCV and MVA-HCV ΔC6L induced similar proportions of HCV-specific CD8^+^ T memory cells, with a predominance of TEM phenotype, which has been described to be required to eliminate the virus through the production of cytolytic molecules and cytokines (62).

To boost efficacy of an HCV vaccine candidate, it should be combined T and B cell responses in a synergistic effect (31), since there is rising evidence that antibodies can be necessary to control HCV replication, especially in early stages of infection (1, 12). A meta-analysis study showed that the inclusion of envelope proteins in a vaccine and therefore the presence of antibodies indicate higher chances of resolving infection in chimpanzees, suggesting that neutralizing antibodies (NAbs) can play a role in protection (8). HCV E2 glycoprotein contains the receptor binding domain and is the major target for Nabs, which interfere with virion’s attachment to cells (63). MVA-HCV and MVA-HCV ΔC6L were able to elicit antibodies against E2 protein in immunized mice, indicating that they not only stimulate T cell response, but also humoral response. Since any effective vaccine should induce both T and B cell immunity in an optimized regimen, our HCV vaccine candidates fulfill these requirements and are able to induce high levels of HCV-specific T cell and humoral immune responses.

Overall, the results show that production of HCV proteins in moDCs infected with MVA-HCV and MVA-HCV ΔC6L modulates expression of host genes involved in innate immunity, decreasing the levels of type I IFN. In infected mice, both MVA-HCV and MVA-HCV ΔC6L induced high, broad and polyfunctional HCV-specific CD8^+^ T cell responses and humoral responses against HCV antigens, making it a good vaccine candidate against HCV. Our results yield valuable information about the use of MVA as a vaccine candidate against HCV that could be tested in future prophylactic and/or therapeutic clinical trials against HCV.

## MATERIALS AND METHODS

### Ethics statement

Animal studies were approved by the Ethical Committee of Animal Experimentation (CEEA) of Centro Nacional de Biotecnología (CNB, Madrid, Spain) in accordance with national and international guidelines and the Royal Decree (RD 53/2013) (permit number PROEX 331/14). Animal experiments were performed at the CNB in a pathogen free barrier animal facility in accordance to the recommendations of the Federation of European Laboratory Animal Science Associations. Buffy coats from healthy blood donors were obtained from the Centro de Transfusion de la Comunidad de Madrid (Madrid, Spain) and their use was approved by their Ethical Committee. Written informed consent was obtained from donors.

### Cells and viruses

DF-1 cells (a spontaneously immortalized chicken embryo fibroblast (CEF) cell line), and primary CEF cells (obtained from specific-pathogen-free 11-day-old eggs; Intervet, Salamanca, Spain) were grown in Dulbecco’s modified Eagle’s medium (DMEM) supplemented with penicillin (100 units/ml; Sigma-Aldrich), streptomycin (100 μg/ml; Sigma-Aldrich), L-glutamine (2 mM; Sigma-Aldrich), nonessential amino acids (0.1 mM; Sigma-Aldrich), gentamicin (50 μg/ml; Sigma-Aldrich), amphotericin B (Fungizone, 0.5 μg/ml; Gibco-Life Technologies), and 10% fetal calf serum (FCS) (Gibco-Life Technologies). The human monocytic THP-1 cell line was cultured in RPMI 1640 medium containing 2 mM L-glutamine, 50 μM 2-mercaptoethanol, 100 units/ml of penicillin, 100 μg/ml of streptomycin (all from Sigma-Aldrich) and 10% FCS. THP-1 cells were differentiated into macrophages by treatment with 0.5 mM phorbol 12-myristate 13-acetate (PMA; Sigma-Aldrich) for 24 h before usage. Freshly isolated peripheral blood mononuclear cells from buffy coats of healthy blood donors were obtained by Ficoll gradient separation on Ficoll-Paque (GE Healthcare), then monocytes were isolated using the Dynabeads Untouched human monocytes kit (Invitrogen) and moDCs were obtained after differentiation of purified monocytes cultivated for 7 days in complete RPMI 1640 medium containing 2 mM L-glutamine, 50 μM 2-mercaptoethanol, 100 units/ml of penicillin, 100 μg/ml of streptomycin (all from Sigma-Aldrich), 10% FCS, 50 ng/ml granulocyte-macrophage colony-stimulating factor and 20 ng/ml interleukin-4 (both from Gibco-Life Technologies). Cells were maintained in a humidified air–5% CO_2_ atmosphere incubator at 37°C.

Poxviruses used in this work included the attenuated MVA-WT, and the recombinant MVA-HCV expressing the nearly full-length HCV genome from genotype 1a (containing Core, E1, E2, p7, NS2, NS3, NS4A, NS4B, NS5A and a part of NS5B), which are inserted into the VACV TK locus of the MVA-WT genome under the transcriptional control of a viral synthetic early/late (sE/L) promoter (26). In addition, MVA-HCV was used as the parental virus for the generation of the recombinant virus MVA-HCV ΔC6L (see later). Virus infections in cells were performed as previously described (64, 65). All viruses were grown in primary CEF cells, purified by centrifugation through two 36% sucrose cushions in 10 mM Tris-HCl (pH 9) and titrated at least two times in DF-1 cells by plaque immunostaining assay, using rabbit polyclonal antibody against VACV strain Western reserve (WR) (Centro Nacional de Biotecnología; diluted 1:1000), followed by an anti-rabbit horseradish peroxidase (HRP)-conjugated secondary antibody (Sigma-Aldrich; diluted 1:1,000), as previously described (66). All viruses were not contaminated with mycoplasma (checked by specific PCR for mycoplasma), bacteria (checked by growth in LB plates without ampicillin) or fungi (checked by growth in Columbia blood agar plates; Oxoid).

### Plasmid transfer vector pGem-RG-C6L wm

The plasmid transfer vector pGem-RG-C6L wm directs the deletion of VACV *C6L* gene from MVA-HCV genome and was used for the generation of the recombinant virus MVA-HCV ΔC6L. pGem-RG-C6L wm was previously generated (22), and contains dsRed2 and rsGFP genes under the control of the sE/L promoter and *C6L* recombination flanking sequences into the plasmid pGem-7Zf (-) (Promega). *C6L* in VACV Copenhagen strain is equivalent to *MVA 019L* in MVA, and for simplicity we used throughout this study the open reading frame nomenclature of Copenhagen strain to refer the MVA genes.

### Construction of MVA-HCV ΔC6L

MVA-HCV ΔC6L deletion mutant generated in this study contains a deletion of the VACV gene *C6L* and was constructed by transfecting plasmid transfer vector pGem-RG-C6L wm into DF-1 cells infected with the parental virus MVA-HCV and then screening for transient Red2/GFP co-expression using dsRed2 and rsGFP genes as transiently selectable markers, as previously described (22, 67). MVA-HCV ΔC6L was finally obtained after six consecutive rounds of plaque purification in DF-1 cells.

### PCR analysis

The correct generation and purity of MVA-HCV ΔC6L was confirmed by PCR with primers spanning the insertion region (VACV TK locus) of the HCV genes and primers spanning VACV *C6L* flanking regions, and by DNA sequence analysis. Thus, viral DNA was extracted from DF-1 cells mock infected or infected at 5 PFU/cell with MVA-WT, MVA-HCV or MVA-HCV ΔC6L, as previously described (26). To verify that deletion of the VACV *C6L* gene was correctly present, primers RFC6L-AatII-F and LFC6L-BamHI-R (previously described in (22)) spanning *C6L* flanking regions were used for PCR analysis of the *C6L* locus. Furthermore, primers TK-L and TK-R (previously described in (26)) annealing in the TK gene-flanking regions, were used for PCR analysis of the TK locus, to verify the correct presence of the HCV genes. All the amplification reactions were performed with Phusion^®^ High-Fidelity DNA Polymerase (New England Biolabs) according to the manufacturer’s recommendations, and the amplification protocol was previously described (67). PCR products were run in 1% agarose gel and visualized by SYBR Safe staining (Invitrogen). The 7,868 Kb HCV DNA insertion and the *C6L* deletion were also confirmed by DNA sequence analysis (Macrogen).

### Expression of HCV proteins

To check the correct expression of HCV proteins by MVA-HCV ΔC6L, monolayers of DF-1 cells were mock infected or infected at 5 PFU/cell with MVA-WT, MVA-HCV or MVA-HCV ΔC6L. At 24 h postinfection (hpi), cells were collected by scraping, centrifuged at 3,000 rpm for 5 min, and cellular pellets were lysed in Laemmli 1X + β-mercaptoethanol. Cell extracts were fractionated by 10% SDS-PAGE and analyzed by Western blotting with mouse polyclonal antibody against Core (C7-50, Thermofisher; diluted 1:5,000), mouse monoclonal antibodies against E1 (Acris Antibodies; diluted 1:1,000), E2 (GenWay Biotech; diluted 1:1,000), and NS5A (GenWay Biotech; diluted 1:1,000) or goat polyclonal antibody against NS3 (Abcam; diluted 1:500), to evaluate the expression of the different HCV proteins. As loading controls, we used rabbit anti-β-actin (Cell Signaling; diluted 1:1,000), and rabbit anti-VACV E3 (Centro Nacional de Biotecnología; diluted 1:1,000) antibodies. An anti-rabbit-horseradish peroxidase (HRP)-conjugated antibody (Sigma; diluted 1:5,000), anti-mouse-HRP-conjugated antibody (Sigma; diluted 1:2,000) or anti-goat-HRP-conjugated antibody (Sigma; diluted 1:5,000) were used as secondary antibodies. The immunocomplexes were detected using an HRP-luminol enhanced chemiluminescence system (ECL Plus; GE Healthcare).

### Analysis of virus growth

To determine the virus growth profile of MVA-HCV ΔC6L, in comparison to that of parental MVA-HCV, monolayers of DF-1 cells grown in 12-well plates were infected in duplicate at 0.01 plaque-forming units (PFU)/cell with MVA-HCV or MVA-HCV ΔC6L. After virus adsorption for 1 h at 37°C, the inoculum was removed, and the infected cells were washed with DMEM without serum and incubated with fresh DMEM containing 2% FCS at 37°C in a 5% CO_2_ atmosphere. At different times postinfection (0, 24, 48, and 72 h), cells were harvested by scraping (lysates at 5 ×; 10^5^ cells/ml), freeze-thawed three times, and briefly sonicated. Virus titers in cell lysates were determined by plaque immunostaining assay in DF-1 cells, as previously described (66).

### RNA analysis by quantitative real-time PCR

Total RNA was isolated using the RNeasy kit (Qiagen) from human THP-1 cells or moDCs mock infected or infected at 5 PFU/cell (for THP-1 cells) or at 0.3 and 1 PFU/cell (for moDCs) with MVA-WT, MVA-HCV or MVA-HCV ΔC6L for 6h. Reverse transcription of 1000 ng of RNA was performed using the QuantiTect Reverse Transcription kit (Qiagen). Quantitative PCR was performed with a 7500 Real-Time PCR System (Applied Biosystems) using the Power SYBR Green PCR Master Mix (Applied Biosystems), as previously described (64). Expression levels of IFN-β, IFIT1, IFIT2, TNF-α and HPRT genes were analyzed by real-time PCR using specific oligonucleotides (their sequences are available upon request). Gene specific expression was expressed relative to the expression of HPRT in arbitrary units (A.U.). All samples were tested in duplicate, and two independent experiments were performed.

### Microarray analysis

Human moDCs from three different healthy donors were mock infected or infected at 1 PFU/cell with MVA-WT, MVA-HCV or MVA-HCV ΔC6L. At 6 hpi, total RNA was isolated with TRIzol (Thermo Fisher) following manufacturer’s recommendations. The three biological replicates were independently hybridized for each transcriptomic comparison. The One-Color Microarray-Based Gene Expression Analysis Protocol (Agilent Technologies, Palo Alto, CA, USA) was used to amplify and label RNA. Briefly, 200 ng of total RNA was reverse transcribed using T7 promotor primer and the cDNA was converted to RNA using T7 RNA polymerase, which simultaneously amplifies target material and incorporates cyanine 3-labeled CTP. 1650 ng of Cy3 probes were mixed and hybridized to a Human Oligo Microarray 4×44K (G2519F_026652, Agilent Technologies) for 17 h at 65°C in Gex Hybridization Buffer HI-RPM in a hybridization oven. Arrays were washed according to the manufacturer’s instructions, dried by centrifugation and scanned at 3 mm resolution on Agilent DNA Microarrays Scanner (G2565BA, Agilent Technologies) and the images were analyzed with Feature Extraction software (Agilent Technologies). Background correction and normalization of expression data were performed using LIMMA (68, 69). LIMMA is part of Bioconductor, an R language project (70). For local background correction and normalization, the methods “normexp” and loess in LIMMA were used, respectively (68). To have similar distribution across arrays and to achieve consistency among arrays, log-ratio values were scaled using as scale estimator the median-absolute-value (68).

Differentially expressed genes were evaluated by the non-parametric algorithm ‘Rank Products’ available as “RankProd” package at Bioconductor (71, 72). This method detects genes that are consistently high ranked in a number of replicated experiments independently of their numerical intensities. The results are provided in the form of p-values defined as the probability that a given gene is ranked in the observed position by chance. The expected false discovery rate was controlled to be less than 5%. Hybridizations and statistical analysis were performed by the Genomics Facility at Centro Nacional de Biotecnología.

For microarray validation, the three donor RNA samples were reverse transcribed and used for real-time PCR to determine the expression levels of IFN-β, IFIT1 and IFIT2 in a similar way as described above.

### Recruitment of immune cells in the peritoneal cavity of C57BL/6 mice

C57BL/6JOlaHsd mice (6 to 8 weeks old) were purchased from Envigo. Groups of mice (n=5) were injected i.p. with 10^7^ PFUs per mouse of MVA-WT, MVA-HCV, MVA-HCV ΔC6L or PBS. At 6, 24 and 48 h post inoculation, peritoneal exudate cells were collected in 6 ml of PBS-2% FCS and the presence of different immune cells was analyzed by flow cytometry, as previously described (73). Briefly, cells were seeded in a 96-well plate and washed twice with PBS Staining (PBS 1X + 0.5% BSA + 1% FCS + 0.065% Sodium Azide + 2 mM EDTA). Then, Fc block (BD Pharmingen) was added to the cells for 15 min in dark at 4°C. For staining the following antibodies were used: CD11c-FITC (clone N418; Bioscience; 1:50), Ly6G-PE (clone 1A8; BD; 1:100), CD11b-PE-Cy7 (clone M1/70; BD; 1:100), F4/80-APC (clone BM8; eBioscience; 1:50), CD19-APC-Cy7 (clone 1D3; BD; 1:100), MHC II (1-A/1-E biotin) (clone 2G9; BD; 1:100) + Avidin-PB (Invitrogen; 1:100), CD45-BV570 (clone 30-F11; Biolegend; 1:100), CD4-PE (clone GK1.5; BD; 1:100), CD8-PE-Cy7 (clone 53-6.7; BD; 1:200), NKp46-APC (clone 29A1.4; Biolegend; 1:100), and CD3-APC-Cy7 (clone 145-2C11; BD; 1:50).

Cells were acquired using a Gallios flow cytometer (Beckman Coulter) and data was analyzed using the FlowJo software (version 8.5.3; Tree Star, Ashland, OR). The different cell populations were gated on CD45^+^ cells as follows: DCs (CD11c^+^/ MHC-II^+^), neutrophils (CD11b^+^/Ly6G^+^), neutrophils α (CD11b^med^/Ly6G^med^), neutrophils β (CD11b^high^/Ly6G^high^), macrophages (F4/80^high^/CD11b^high^), B cells (CD19^+^), B cells High (CD19^+^/CD11b^+^), CD8 T cells (CD3^+^/CD8^+^), CD4 T cells (CD3^+^/CD4^+^), NKs (NKp46^+^/CD3^−^) and NKT cells (NKp46^+^/CD3^+^). The absolute numbers of immune cell populations for each mouse were determined by flow cytometry after extrapolation to the number of cells counted after the peritoneal washes.

### Peptides

Purified HCV peptide pools of the HCV virus H77 strain (genotype 1a), were obtained through BEI Resources (National Institute of Allergy and Infectious Disease, National Institutes of Health). Peptides cover the entire HCV H77 genome as consecutive 13-to 19-mers overlapping by 11 or 12 amino acids. Peptides were resuspended to a final concentration of 1 mg/ml and grouped into seven pools: Core pool (28 peptides), E1 pool (28 peptides), E2 pool (55 peptides), p7 + NS2 pool (40 peptides), NS3 pool: comprising NS3-1 (49 peptides) plus NS3-2 (49 peptides), NS4 pool (47 peptides), and NS5 pool: comprising NS5-1 (55 peptides) plus NS5-2 (53 peptides), and NS5-3 (53 peptides). Peptides were used for *ex vivo* stimulation of splenocytes from immunized mice.

### C57BL/6 mice immunization schedule

Female C57BL/6JOlaHsd mice (6 to 8 weeks old) were acquired from Envigo. An MVA prime/MVA boost immunization protocol was performed to study the immunogenicity of MVA-HCV and MVA-HCV ΔC6L vaccine candidates. Mice of each group (n=8) were immunized with 10^7^ PFU/mouse of MVA-HCV, MVA-HCV ΔC6L or MVA-WT by i.p. route in 200 μl of PBS and 2 weeks later received a second dose of 10^7^ PFU/mouse of the same virus as in the prime. At 10 and 53 days after the last immunization, 4 mice in each group were sacrificed and spleens were processed to measure by ICS assay the adaptive and memory HCV-specific T cell immune responses, respectively. Furthermore, serum samples were obtained at the same time points to analyze the HCV-specific humoral immune responses. Four independent experiments were performed.

### ICS assay

The magnitude, breath, polyfunctionality, and phenotype of the HCV-specific T cell responses were analyzed by ICS, as previously described (22, 24, 26, 67, 74). After spleen disruption, 4 ×; 10^6^ splenocytes were treated with NH_4_Cl 0.1M in ice for 5 min to deplete the red blood cells. Then, splenocytes were seeded on 96-well plates and stimulated for 6 h in complete RPMI 1640 medium supplemented with 10% FCS containing 1 µl/ml of GolgiPlug (BD Biosciences), monensin 1X (eBioscience), anti-mouse CD107a-Alexa Fluor^®^ 488 (clone 1D4B; eBioscience; 1:300), and 1 µg/ml of the different HCV peptide pools (Core, E1, E2, p7+NS2, NS3, NS4, and NS5). Then, cells were washed with PBS 1X-2% FCS-2 mM EDTA (FACS Buffer), stained using 0.5 µl/ml of LIVE/DEAD™ Fixable Violet Dead Cell Stain Kit (Invitrogen) for 30 min at 4°C and washed once with FACS Buffer. Cells were then stained for 20 min at 4°C in FACS buffer containing the following fluorochrome-conjugated antibodies against different cell surface markers: PE-CF594 hamster anti-mouse CD3e (clone 145-2C11; BD; 1:100), APC-Cy™7 rat anti-mouse CD4 (clone GK1.5; BD; 1:100), V500 rat anti-mouse CD8a (clone 53-6.7; BD; 1:100), Alexa Fluor^®^ 700 rat anti-mouse CD62L (clone MEL-14; BD; 1:100), and PerCP-Cy5.5 rat anti-mouse CD127 (clone A7R34; eBioscience; 1:100). After one wash the cells were fixed and permeabilized for 20 min at 4°C using the Cytofix/Cytoperm™ kit (BD), following by two washes in perm/wash buffer and addition of purified rat anti-mouse CD16/CD32 (Mouse Fc block™; clone 2.4G2; BD; 1:100) for 15 min at 4°C. Next, cells were stained intracellularly for 20 min at 4°C in perm/wash buffer containing the following fluorochrome-conjugated antibodies: PE-Cy™7 rat anti-mouse IFN-γ (clone XMG1.2; BD; 1:100), PE rat anti-mouse TNFα (clone MP6-XT22; eBioscience; 1:100), and APC rat anti-mouse IL-2 (clone JES6-5H4; BD; 1:40). Following staining, cells were washed twice in perm/wash buffer and finally resuspended in FACS buffer. Furthermore, cells that were stained individually with each fluorochrome-conjugated antibody were used as controls for compensation. Cells were acquired using a Gallios flow cytometer (Beckman Coulter) and flow cytometry data was analyzed using FlowJo software (version 8.5.3; Tree Star, Ashland, OR). After gating every cytokine or surface marker, boolean combinations of single functional gates were then created to determine the frequency of each response based on all possible combinations of cytokine expression or all possible combinations of differentiation marker expression. Background responses detected in negative-control samples (stimulated with RPMI) were subtracted from those detected in stimulated samples for every specific functional combination.

### ELISA

Total IgG levels of binding antibodies to HCV E2 protein present in the sera of immunized mice were determined using ELISA. Ninety-six-well plates (Nunc MaxiSorp^®^) were coated with 50 µl of purified recombinant HCV (genotype 1a, isolate H77) E2 protein (Sino Biological) at a concentration of 2 µg/ml at 4°C overnight. Plates were then washed 3 times with PBS 1X supplemented with 0.05% Tween-20 and blocked with 200 µl of PBS-5% milk-0.05% Tween-20 for 2 h at room temperature (RT). Next, serum samples were diluted 1:100 in PBS-1% milk-0.1% Tween-20 and 100 µl were added to the plates following an incubation of 1.5 h at RT. Then, plates were washed three times with PBS-0.05% Tween-20 and 100 µl of secondary HRP-conjugated goat anti-mouse IgG antibody (SouthernBiotech, AL, USA, diluted 1:1,000 in PBS-1% milk-0. 1% Tween-20) was added and incubated for 1 h at RT. After incubation, plates were washed 3 times with PBS-0.05% Tween-20 and 100 µl of 3,3′,5,5′-tetramethylbenzidine (TMB) (Life Technologies) were added. After 15 min 50 µl of H_2_SO_4_ 1M were added to stop the colour development. Absorbance was read at 450 nm.

### Statistical procedures

Significant differences (^*^ =P ≤ 0.05; ^**^ =P ≤ 0.01; ^***^ = P ≤ 0.001) in RT-PCR data were calculated using the Holm-Sidak method, with alpha=5%. Differentially expressed genes in microarray analysis were selected after application of the following statistical filters (FDR Prod P value <0.05 or P value LIMMA < 0.05 and fold change ≥ 2 or ≤ −2). The statistical significance of differences between groups in the experiment of cell recruitment was determined by Student’s t test (unpaired, two tailed). Statistical analysis of the ICS data was realized as previously described (67, 75) using an approach that corrects measurements for the negative-control medium response (RPMI), with calculation of confidence intervals and P values. Only antigen response values significantly larger than the corresponding RPMI values are represented. Background values were subtracted from all of the values used to allow analysis of proportionate representation of responses.

## ACKNOWLEDGMENTS

This research was supported by Spanish grant SAF-2013-45232-R and SAF-2017-88089-R. We thank Cristina Sánchez Corzo and Victoria Jiménez for expert technical assistance in cell culture, virus growth and purification. We also thank to Gloria García Casado at the Genomics Facility at Centro Nacional de Biotecnología for their valuable help in the microarray analysis. María Q. Marín received a Formación del Profesorado Universitario PhD fellowship, from the Spanish Ministry of Education and a short-term EMBO fellowship.

